# Genetic determinants of multi-drug resistance in *Acinetobacter baumannii* at an academic hospital in Pretoria, South Africa

**DOI:** 10.1101/809129

**Authors:** Noel-David Nogbou, Dikwata Thabiso Phofa, Maphoshane Nchabeleng, Andrew Munyalo Musyoki

**Affiliations:** Department of Microbiological Pathology, Sefako Makgatho Health Sciences University, Pretoria, South Africa; National Health Laboratory Service, Dr George Mukhari Tertiary Laboratory, Pretoria, South Africa

## Abstract

Antimicrobial resistance is now globally recognised as the greatest threat to human health. *Acinetobacter baumanniis’* (*A. baumannii*) clinical significance has been driven by its ability to obtain and transmit antimicrobial resistance factors. In South Africa, *A. baumannii* is a leading cause of healthcare associated infections (HAI). In this study, we investigated the genetic determinants of multi-drug resistant *A. baumannii* (MDRAB) at a teaching hospital in Pretoria, South Africa.

One hundred non repetitive isolates of *A. baumannii* were collected for the study at Dr George Mukhari Tertiary Laboratory (DGMTL). Antimicrobial susceptibility testing was performed using the VITEK2 system (bioMerieux, France). The prevalence of common resistance associated genes and AdeABC efflux pump system associated genes were investigated using conventional PCR. Genetic relatedness of isolates was then determined using rep-PCR.

Seventy (70) of 100 isolates collected were confirmed to be multi-drug resistant and were *bla*_*OXA51*_ positive. Phenotypically, the isolates where resistant to almost all tested antibiotics. However, one isolate showed intermediate susceptibility to tigecycline while all were susceptible to colistin. Oxacillinase encoding gene *bla*_*OXA-23*_ was the most detected at 99% and only 1% was positive for *bla*_*OXA-40*_. The PCR results for metallo-betalactamase (MBL) encoding genes showed that MBL *bla*_*VIM*_ was the most frequently detected at 86% and *bla*_*SIM-1*_ at 3% was the least detected. Out of 70 isolates, 56 isolates had the required gene combination for an active efflux pump. The most prevalent clone was clone A at 69% of the isolates. Regarding treatment; colistin and tigecycline are the most effective against strains encountered at DGMTL as all tested carbapenems seem to have lost their effectiveness.

The major genotypic determinants for drug resistances are oxacillinases: *bla*_*OXA-51*_ (100%) and *bla*_*OXA-23*_ (99%). The study reports for the first time, *bla*_*OXA-40*_ and *bla*_*SIM-1*_ detection in *A. baumannii* in South Africa.

## Background

Antibiotics are at the forefront of the battle against bacteria and other potentially dangerous infectious agents to human health. The use of antibiotics has changed the outcome of bacterial infections and saved millions of lives [1]. On the 27^th^ of February 2017 in Geneva, World Health Organisation (WHO) released a list of bacteria that have become resistant to multiple classes of antibiotics including carbapenems and third-generation cephalosporins [2]. Multi-drug resistant *Acinetobacter baumannii* (MDRAB) has been identified as one of the leading clinically relevant multi-drug resistant organism that threatens human health [2]. In South Africa, *A. baumannii* is the leading cause of nosocomial infections [3] and its prevalence of isolation in various clinical samples has been increasing over time [4; 5; 6]. MDRAB’s clinical importance has been driven by its ability to obtain and transmit antimicrobial resistance genes [5] and to cause outbreaks in health care settings [7]. It is associated with high morbidity, high mortality, prolonged hospitalization and increased cost of hospitalisation [7; 3; 8].

Resistance genes are acquired through various mechanisms and enabling A. baumannii to resist action of several antibiotic families. Combination of mechanisms such as an increased expression of oxacillinase (OXA)-type carbapenemases and non-enzymatic mechanisms, such as decreased cell membrane permeability, overexpression of multi-drug efflux pump proteins and/or alterations in penicillin-binding proteins are reported to induce multiple drug resistance in *A. baumannii* [9]. For effective infection control and treatment, it is important to investigate and report on the diversity of prevalent strains. In this study, genetic determinant of MDRAB at Dr George Mukhari Tertiary Laboratory (DGMTL) were investigated.

## Materials and methods

### Sample collection

Ethical clearance to conduct the study was granted by Sefako Makgatho Health Sciences University Research and Ethics Committee. A hundred non repetitive isolates of MDRAB identified by VITEK2 system (bioMerieux, France) were collected at Dr George Mukhari Tertiary Laboratory (DGMTL) located at Dr George Mukhari Academic Hospital; from March 2017 to August 2017 and February 2018 to April 2018. Strains of *A. baumannii* were considered multi-drug resistant when resistant to at least one antibiotic in three different antibiotic classes [10].

### Antimicrobial susceptibility testing

Antimicrobial susceptibility of isolates was established using the VITEK2 system (bioMerieux, France). Piperacillin + tazobactam, ceftazidime, cefepime, cefotaxime/ceftriaxone, imipenem, meropenem, trimethoprim/sulfamethoxazole, gentamycin, ciprofloxacin, tigecycline and colistin were tested.

### Confirming multi-drug resistance and genotypic identification of *A. baumannii* isolates

Multi-drug resistance profile was confirmed using the disk diffusion method according to Machanda *et al.*, [10]. A standardized inoculum of the isolate was prepared using normal saline (Diagnostic media product (DMP), NHLS, South Africa) and adjusted to 0.5 McFarland using a turbidity meter (DensiCHEK plus bioMerieux, France). The suspension was lawned on to Muller Hinton (MH) (Diagnostic media product, NHLS, South Africa (DMP)) agar plate. Susceptibility to one of each of the different classes of antibiotics was tested as follow: Gentamicin 10μg for aminoglycosides; Ciprofloxacin 5μg for quinolones and colistin 10μg for polypeptides. For beta-lactams, Piperacillin + Tazobactam 110μg for penicillins combined with inhibitors, ceftriaxone 30μg and ceftazidime 30μg for cephalosporins, meropenem 10μg for carbapenems were used. Antibiotic disks were placed onto the inoculated agar plate. The process was carried out according to CLSI guidelines (2017; CLSI Document M100-S27). The OXA-51 is a naturally occurring oxacillinase in *A. baumannii* that can be used to reliably identify this organism genotypically [11; 12; 13]. Therefore, it was used to confirm the genotypic identification of the *A. baumannii* isolates in this study in addition to the phenotypic methods.

### Determining the prevalence of common antimicrobial resistance genes

#### DNA extraction

DNA extraction was performed using the boiling method as described by Olive and Bean [14] with a slight modification. Briefly, from each fresh overnight culture, a loopful of bacteria was taken from MH agar plates and suspended in 1mL of saline (Normale Saline G121721, DMP; South Africa) in an eppendorf tube, then centrifuged (MIKRO 200, Labotec, South Africa) at 7500rpm for 30 minutes. The supernatant was discarded and the pellet was re-suspended in 200μL of PCR water (Water for Molecular Biology, BioConcept Ltd, Switzerland). The obtained suspension was centrifuged at 7500rpm for 20 minutes. Thereafter, the suspension was boiled at 95 °C for 20 minutes using a thermomixer (Thermomixer Compact Eppendorf 5350 MERCK Chemical Pty. Ltd, South Africa), then centrifuged for 5 minutes at 7500rpm. The supernatant was used as the template for PCR immediately or stored at −20 °C until use.

#### Master Mix preparation

PCR master mix was prepared by using MyTaqTM HS ready mix (Bioline, UK) to detect genes of interest in *A. baumannii*. All PCR primers used in this study were synthesized by a commercial vendor in South Africa. Primers are listed in Supplimentary1 Table (S1). The Bioline protocol (Bioline; UK) was followed to prepare multiplex PCR assays using primer pairs with similar melting temperatures and monoplex PCR for primer pairs with different melting temperatures. PCR was performed in a reaction mixture of a total volume of 25μL; 12.5μL of My Taq ™ Red Mix (Bioline; UK), 0.5μL of each primer (forward and backward) and PCR grade water (Water for Molecular Biology, BioConcept Ltd, Switzerland) was added to make up to 20μL and 5μL of DNA template was added to constitute a 25μL reaction mix. The thermocycling conditions used for the detection of drug resistance associated genes are listed in supplementary 2 Table (S2).

### Determining genetic relatedness of *A. baumannii* isolates

The genetic relatedness was determined using repetitive extragenic palindromic PCR (rep-PCR). Samples that were confirmed to be MDR by VITEK2 and *bla*_*OXA-51*_ positive by PCR underwent rep-PCR. The REP regions were amplified according to Santimaleeworagun *et al.*, [15] using MyTaqTM HS mix (Bioline, UK). Briefly, a total volume of 25μL was prepared as mastermix with 5μL as DNA template and 20μL constituting of My Taq ™ Red Mix (Bioline; UK) 12.5μL; 0.5μL of each specific primer (forward and reverse) and adjusted to 20μL using PCR water (Water for Molecular Biology, BioConcept Ltd, Switzerland). Thermocycling conditions are listed in S2 Table.

### PCR amplicon detection

PCR amplicons were separated and detected using ethidium bromide stained agarose gel electrophoresis and visualized under UV light. The expected amplicon sizes for the various primer sets and rep-PCR oligonucleotides are indicated in S1 Table.

### Gene burden and statistical analysis

The gene burden of overall 11 targeted genes was established using IBM SPSS Statistics 25 software. Statistical analysis with 95% confidence level and p-value was assessed using IBM SPSS Statistics 25 software.

### Summary of research methodology

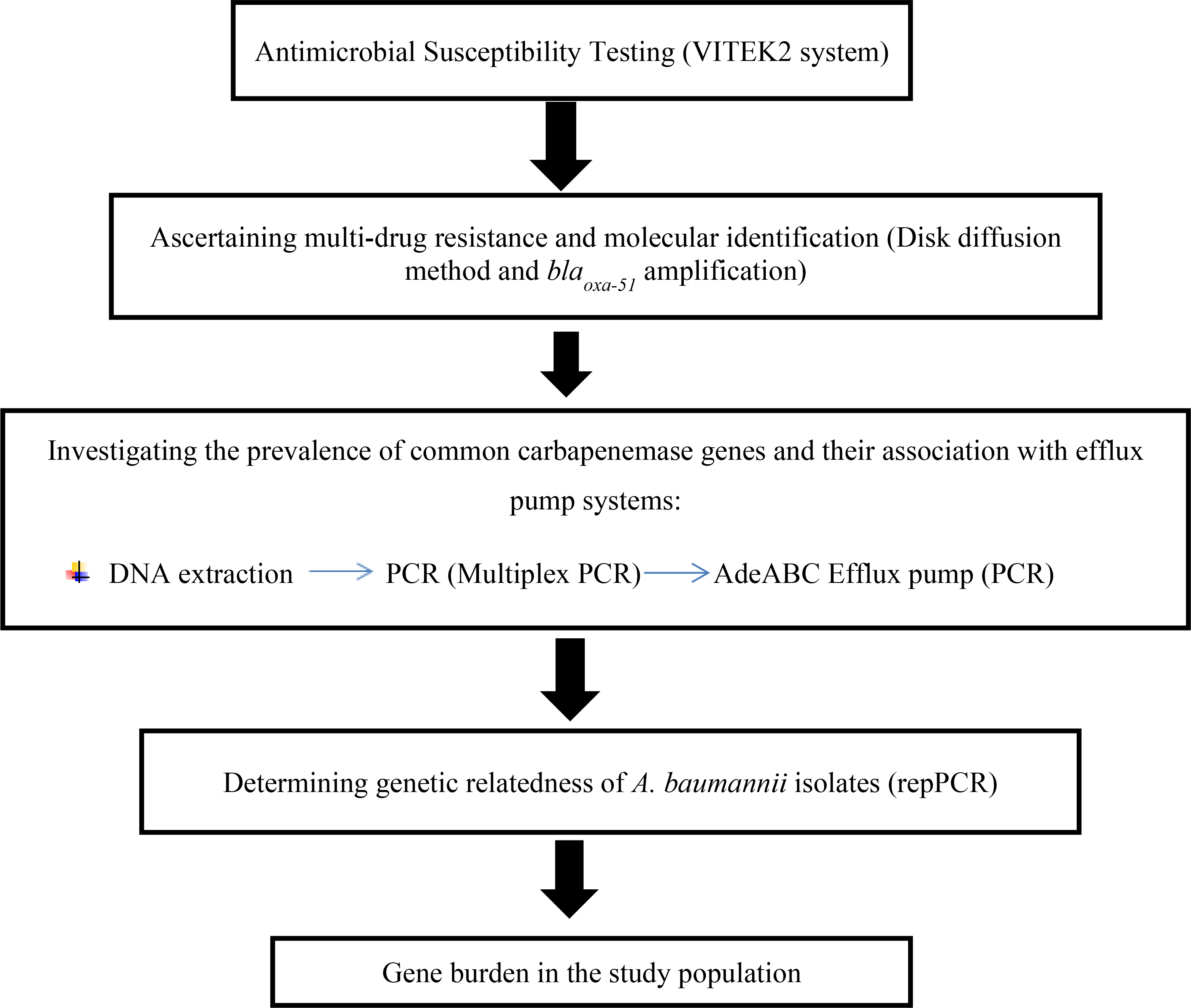

## Results

### Antimicrobial susceptibility testing

Eleven antibiotics were tested for susceptibility on all 100 study isolates. All isolates (100%) were resistant to ceftazidime, cefepime and piperacillin + tazobactam; 98% of the isolates were resistant to trimethoprim/sulfamethoxazole and cefotaxime/ceftriaxone; 95% of the isolates were resistant to imipenem and meropenem; 90% of isolates were resistant to gentamycin; 88% of isolates were resistant to ciprofloxacin; 1 isolate showed reduced susceptibility to tigecycline with the rest (99%) of the isolates being susceptible. All the isolates (100%) were susceptible to colistin (Table 1).

**Table 1.**
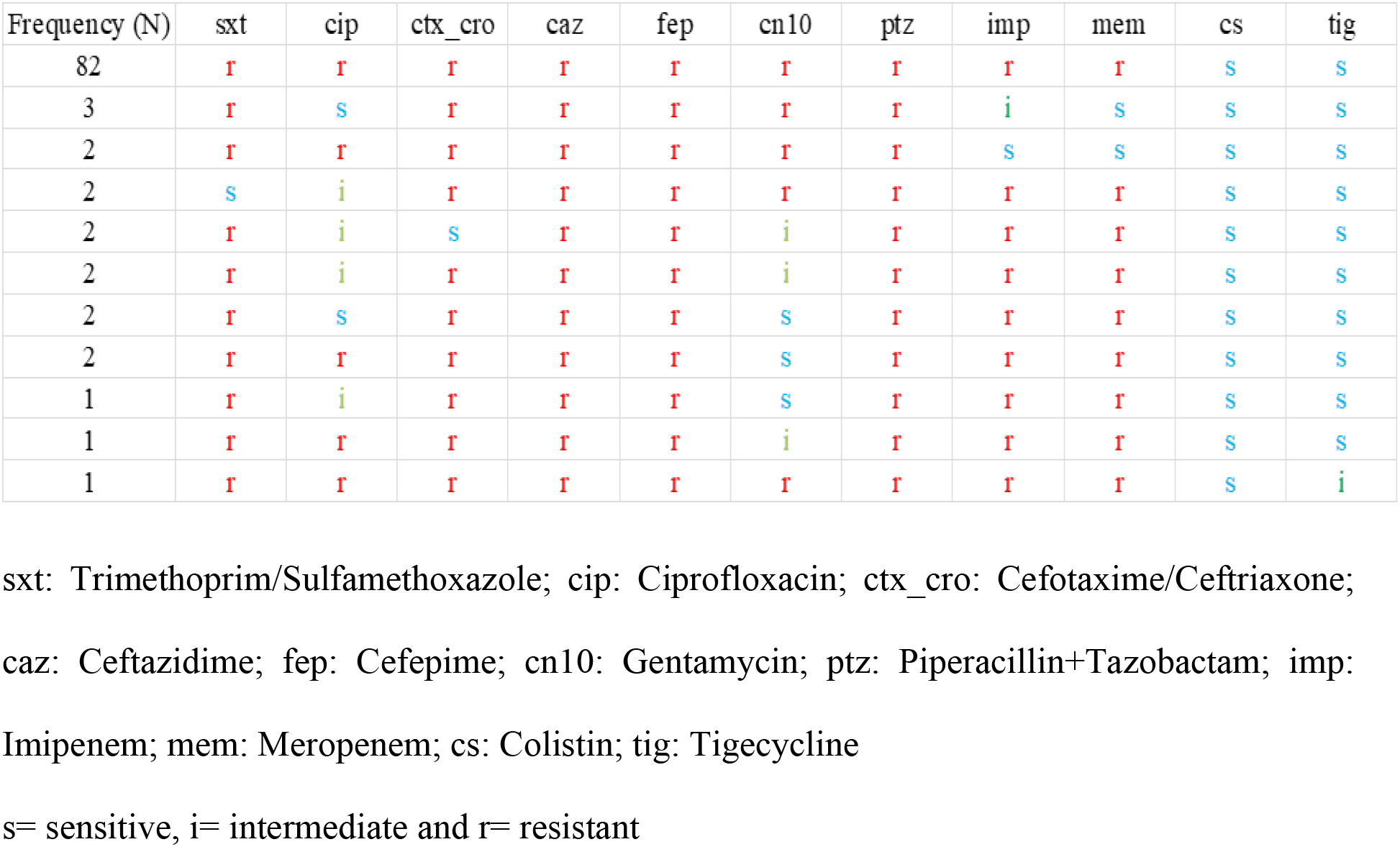
Summary of the VITEK2 system results.

### Confirming multi-drug resistance and molecular identification of *A. baumannii* isolates

Seventy (70) of 100 isolates collected were confirmed *A. baumannii* on the basis of the presence of *bla*_*OXA-51*_ genes and confirmed multi-drug resistant based on Manchanda *et al.,* [10] criteria. These isolates constituted the final study samples. The 30 remaining isolates were excluded from our study; 17 of these were excluded because they were not multi-drug resistant; 9 isolates were not multi-drug resistant and were negative for *bla*_*OXA-51*_ and 4 of them were multi-drug resistant but *bla*_*OXA-51*_ negative.

### Determining the prevalence of common resistance associated genes

From the 70 study samples, oxacillinase encoding gene *bla*_*OXA-23*_ was the most frequently detected with 69 (98%) positive isolates and only 1 (1%) positive isolate for each *bla*_*OXA-58*_ and *bla*_*OXA-40*_ Table 2. PCR conducted for metallo-betalactamase (MBL) encoding genes detected 60 (86%) *bla*_*VIM*_ and 41 (58%) *bla*_*NDM*_ positive isolates, followed by 5 (7%) *bla*_*IMP*_ and 2 (3%) *bla*_*SIM-1*_ with positive isolates Table 2. Fifty-six out of 70 isolates tested positive for the required gene combination for an active efflux pump. Nine isolates had the structural gene with no regulatory genes while 2 isolates had the structural gene and only one of the regulatory genes (*AdeS*). Three isolates did not have any of the 3 genes.

**Table 2.** Summary of PCR results.

| CLASS OF ENZYMES | TARGET GENES | POSITIVE FREQUENCY n=70 (%) |
| --- | --- | --- |
| **Oxacillinases** | *bla<sub>OXA-51</sub>* | 70 (100%) |
|  | *bla<sub>OXA-23</sub>* | 69 (98%) |
|  | *bla<sub>OXA-40</sub>* | 1 (1%) |
|  | *bla<sub>OXA-58</sub>* | 1 (1%) |
| **Metallobeta-lactamases (MBL)** | *bla<sub>VIM</sub>* | 60 (86%) |
|  | *bla<sub>IMP</sub>* | 5 (7%) |
|  | *bla<sub>NDM</sub>* | 41 (58%) |
|  | *bla<sub>SIM-1</sub>* | 2 (3%) |
| **Efflux pump resistance associated genes** | *adeb* | 67 (96%) |
|  | *adeR* | 56 (80%) |
|  | *adeS* | 58 (83%) |

### Gene burden in the study population

Of the 11 investigated genes; 1 isolate was positive for 10 genes and another isolate for 8 genes. Eight (8) isolates were positive for 7 genes; 14 isolates were positive for 6 genes; 24 isolates were positive for 5 genes; 7 isolates were positive for 4 genes; 12 isolates were positives for 3 genes and 3 isolates were positive for 2 genes.

### Determining genetic relatedness of *A. baumannii* isolates

The strains of MDR *A. baumannii* were grouped according to band patterns as described by Santimaleeworagun *et al.,* [15]. The most common clone was clone A with 49 (69%) isolates, followed by clone B and D with 8 (12%) each. Clone C and E were less represented with 2 (3%) and 3 (4%) isolates respectively (Fig 1).

**Fig 1.**
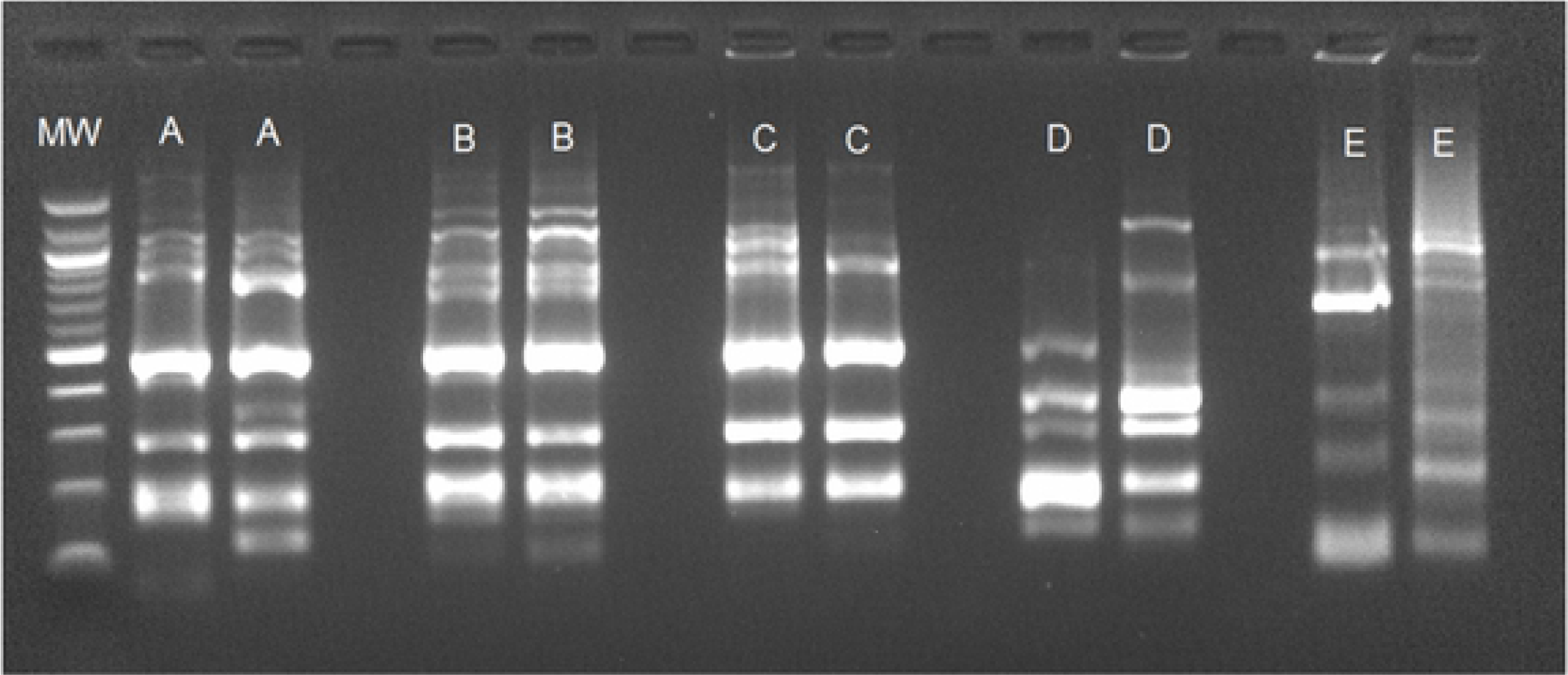
Genetic relatedness of study isolates. **MW**: Molecular Weight. **Clone**: A, B, C, D and E

## Discussion

### Antimicrobial susceptibility testing

Bacteria have developed several resistance mechanisms against antibiotics and are still developing new ways to overcome novel molecules [16]. In this study,100 isolates identified by VITEK2 system as MDRAB reveiled a wide range of resistance to various antibiotics. Colistin was the only agent showing susceptibility among all the study isolates. Only 1 isolates showed intermediate susceptibility to tigecycline (Table 1). Similar results were observed by Lowings *et al.,* [6], during a study conducted in Pretoria, South Africa. The study reported resistance to imipenem at 86%; meropenem at 86%; cefepime at 90% and ceftazidime at 89%. Our study results revealed higher resistance prevalence compared to a similar study by Kock *et al.,* [5]. In their study also conducted in Pretoria (2008), they reported resistance to imipenem at 59 %; meropenem at 63 %; cefepime at 62 % and ceftazidime at 45 %. The prevalence of drug resistance to antimicrobials other than the polymixin class in *A. baumannii* isolates has increased to 100% over the years [5; 17]. To date, Colistin has remained susceptible in all the *A. baumannii* isolates collected in this region. Our study noted with concern; the occurrence of intermediate susceptibility to Tigecycline, which is the only other drug with activity against MDRAB isolates in this study. The ability of *A. baumannii* to acquire and transmit drug resistance genes through several mechanisms such as transfer of integrons, plasmids or transposons and mutation of endogenous genes is associated with among others poor infection control practices in health care settings [18; 5].

### Confirming multi-drug resistance and identification of *A. baumannii* isolates

Thirteen percent (13%) of the isolates were not *A. baumannii* and 87 (87%) out of 100 isolates were confirmed *A. baumannii* after detection and amplification of *bla*_*OXA-51*_ Table 2. Our results are similar to those reported by Kock *et al.,* [5]. In their study, 19% of isolates identified by VITEK2 as *A. baumannii* were negative for *bla*_*OXA-51*_. Consequently, they were considered as misidentified. The oxacillinase *bla*_*OXA-51*_ has been used as a simple and reliable genotypic way of identifying *A. baumannii* strains [12]. A similar study by Lowings *et al.,* [6] reported that 99% *A. baumannii* isolates collected were positive for *bla*_*OXA-51*_ and only 1% of isolate was negative for *bla*_*OXA-51*_. In that study, isolates were identified using an alternative method (MALDI TOF-MS) which is known to have higher sensitivity than VITEK2. Six (6) isolates were found to be misidentified as *A. baumannii* by VITEK2 system. Therefore, our identification method may have included other species outside of the *A. baumannii* complex. This may explain the 13 (13%) *bla*_*OXA-51*_ PCR negative isolates. Among the 87 (87%) confirmed *A. baumannii* isolates, 17 were not multi-drug resistant according to the definition criteria proposed by Manchanda *et al.,* [10]. Consequently, they were removed from further analysis.

### Determining the prevalence of common drug resistance associated genes

A high prevalence of oxacillinase encoding gene *bla*_*OXA-23*_ was noted in 69 (99%) of the 70 isolates Table 2. A study conducted in Thailand also detected the *bla*_*OXA-23*_ in 42 out of 43 (99%) *A. baumannii* isolates [15]. A similar report by Liakopoulos *et al.,* [19] in Greece, noted that 120 out of 127 (95 %) *A. baumannii* isolates collected in 2011 were *bla*_*OXA-23*_ positive. Oxacillinase encoding gene *bla*_*OXA-23*_ was also reported as the most common carbapenamase encoding gene among *Acinetobacter spp* by Corrêa *et al.,* [20] in Brazil and Santimaleeworagun *et al.,* [15] in Thailand. It has been associated with resistance to carbapenems [21]. This study reported resistance rates of 95% to imipenem and meropenem (Table 1). Our results show higher resistance rates than previously reported in the region: 77% by Lowings *et al.,* in Tshwane region and 59% by Kock *et al.,*in Pretoria [5]. Clearly, the prevalence of *bla*_*OXA-23*_ in *A. baumannii* isolates in Pretoria is increasing. This observation is not unique to the Pretoria region. Globally there is an increase in OXA-23-producing *A. baumannii* [22; 23; 24]. This increased occurrence can be due to the acquisition of mobile genetic elements [25] and/or poor infection control practices [26].

In this study, only 1% of the isolates had the oxacillinase encoding gene *bla*_*OXA-58*_ and *bla*_*OXA-40*_ Table 2. A study conducted on *A. baumannii* isolates by Lowe *et al.,* [17] in South Africa on 69 isolates, reported the *bla*_*OXA-58*_ gene in 4% of their study samples. In 2015, the same team tested 100 isolates for the *bla*_*OXA-58*_ gene and found no positive isolates [6]. A study in Thailand isolated only 1 sample with *bla*_*OXA-40*_ out of 43 isolates [15]. These genes were found to be low in our area during our study. The *bla*_*OXA-58*_ and *bla*_*OXA-40*_ are also associated with resistance to carbapenems [27; 28]. However, strict infection prevention and control measures are needed to avoid an increase in the prevalence of isolates harbouring these genes.

Of the 70 isolates, the most frequently detected MBL genes were: *bla*_*VIM*_ and *bla*_*NDM*_ at 86% and 58% respectively Table 2; *bla*_*IMP*_ and *bla*_*SIM*_ were detected in 7% and 3% of the isolates respectively Table 2. In South Africa, since their first report in 2008 [5]; 2011 [29] and 2012 [30]; the occurrence of *bla*_*VIM*_, *bla*_*NDM*_ and *bla*_*IMP*_ respectively has increased over the years [31]. MBLs have mostly been reported in carbapenemase-producing Enterobacteriaceae [32; 31]. Thus, the ability of *A. baumannii* to easily acquire resistant genes and their high prevalence in hospital settings may explain the increasing prevalence of these genes [33; 25]. These results are consistent with the national picture on the progression of MBL producing strains. Two isolates (3%) were positive for *bla*_*SIM-1*_. MBL *bla*_*SIM-1*_ is rarely found in *A. baumannii* [34]. This is the first report on *bla*_*SIM-1*_ in South Africa. The overall increased occurrence of MBL suggests rapid dissemination of these genes once introduced into the environment.

Overexpression of efflux pumps is known to be an *A. baumannii* virulence factor and is associated with decreased susceptibility to several antimicrobial agents [35]. The *adeABC* operon encodes AdeA membrane fusion protein, multi-drug transporter protein AdeB and AdeC outer membrane protein. The AdeR-AdeS; a two-component system, are genes that regulate *adeABC* operon [36]. It has been reported that constitutive overexpression of the adeABC efflux system is due to either upstream insertion of IS*aba-1* on *adeABC* operon or by a single or multiple-points mutation (SNP) in the *adeR* and *adeS* genes [37]. These two conditions induce a decreased intracellular concentration of antimicrobials leading to resistance not only to fluoroquinolones but also aminoglycosides, tetracyclines,
chloramphenicol and beta-lactams [38]. In a study conducted on 50 *A. baumannii* isolates by Lari *et al.,* 16 (32%) isolates were reported to have an active efflux pump after phenotypic testing [37]. Higgings *et al.,* [38] suggested that an increased *adeB* induced resistance to ciprofloxacin. In contrast, Bratu *et al.,* [39] suggested that the increased *adeB* expression by itself did not play a major role in fluoroquinolone resistance and that additional mechanisms should be investigated. In our study, the suitable combination (*AdeB*+; *AdeS*+ and *AdeR*+) for a potential active efflux pump was present in 80% of the study isolates. This correlates with the phenotypic results reported by VITEK2. Though the *ISaba-1* insertion or SNPs were not investigated in our study, the correlation between potential active efflux pumps and resistance to aminoglycosides; quinolones; cephalosporins and beta-lactams (Table 1 and Table 2) supports the suggestion that active efflux pumps may play a role in the antimicrobial resistance in *A. baumannii* isolates collected at DGMTL.

### Determining genetic relatedness of *A. baumannii* isolates

Rep-PCR differentiates bacterial strains by using primers complementary to interspersed repetitive consensus sequences that enable amplification of diverse-sized DNA fragments consisting of sequences between the repetitive elements. Understanding strain diversity is important in assessing progress and outcomes of infection prevention and control measures. In this study, clone A was the most prevalent (Fig 1). However, 4 others lineages were found to co-exist in our setting. A study conducted in a burn unit in the university hospital of Chicago demonstrated an *Acinetobacter calcoaceticus* complex outbreak among patients that lasted 21 months [43]. This suggests that infection prevention and control (IPC) measures should be reinforced. An efficient IPC program may minimize the reservoir for bacterial transmission in hospital settings [44].

### Gene burden in the study population

The presence of specific genes in isolates is associated with a specific mechanism of resistance. This exercise gives us a picture of an increasing number of strains of bacteria able to express several genes responsible for various resistance mechanisms. Since the 1970s antimicrobial resistance genes in *A. baumannii* have progressively increased. By 2007, up to 70% of isolates in certain settings were multi-drug resistant [40]. The increasing occurrence of antimicrobial resistance genes in *A. baumannii* strains reduces treatment options [17; 41]. In this study, 63 over 70 isolates had more than 3 antimicrobial resistance genes. This is consistent with the increase of resistance globally [3; 42] and MDRAB specifically in our hospital setting.

### Study limitation

Despite the findings, this study carries some limitations. The demographic data collected did not include the residential area of patients from whom the samples were collected; this limits extrapolation of results to reflect the general population in South Africa. The study only used a genetic fingerprinting method that is relevant for establishing relatedness locally; a global established genetic typing method would improve the global relevance of strain typing for this study.

## Conclusion

After evaluating 100 *A. baumannii* isolates, the study concluded that isolates collected at DGMTL are multi-drug resistant due to various mechanisms. The major genotypic determinants for drug resistances were for oxacillinases: *bla*_*OXA-51*_ (100%), *bla*_*OXA-23*_ (98%). We report the first *bla*_*OXA-40*_ and *bla*_*SIM-1*_ positive *A. baumannii* isolates in South Africa. Colistin and tigecycline are the most active antimicrobial agents against *A. baumannii* isolates encountered at DGMTL. Judicious prescription, routine surveillance to promote early detection of resistance to commonly used antibiotics are needed in order to forestall the menace of multiple drug resistant *A. baumannii* isolates.

## Acknowledgements

We would like to acknowledge the support by SMU research development grant and the NHLS staff that assisted in one way or the other during this study.

## Supporting information

**S1 Table. The oligonucleotides sequences for PCR**

**S2 Table. Thermocycling condition used in the study**

## References

1. Rossolini GM, Arena F, Pecile P, Pollini S. Update on the antibiotic resistance crisis. Current opinion in pharmacology. 2014 Oct 1;18:56–60. doi: 10.1016/j.coph.2014.09.006

2. World Health Organization. WHO publishes list of bacteria for which new antibiotics are urgently needed. 2017.

3. Gounden R, Bamford C, van Zyl-Smit R, Cohen K, Maartens G. Safety and effectiveness of colistin compared with tobramycin for multi-drug resistant *Acinetobacter baumannii* infections. BMC infectious diseases. 2009 Dec;9(1):26. doi: 10.1186/1471-2334-9-26

4. Ahmed NH, Baba K, Clay C, Lekalakala R, Hoosen AA. In vitro activity of tigecycline against clinical isolates of carbapenem resistant *Acinetobacter baumannii* complex in Pretoria, South Africa. BMC research notes. 2012 Dec;5(1):215. doi: 10.1186/1756-0500-5-215

5. Kock MM, Bellomo AN, Storm N, Ehlers MM. Prevalence of carbapenem resistance genes in *Acinetobacter baumannii* isolated from clinical specimens obtained from an academic hospital in South Africa. Southern African Journal of Epidemiology and Infection. 2013 Jan 1;28(1):28–32. https://doi.org/10.1080/10158782.2013.11441516

6. Lowings M, Ehlers MM, Dreyer AW, Kock MM. High prevalence of oxacillinases in clinical multidrug-resistant *Acinetobacter baumannii* isolates from the Tshwane region, South Africa–an update. BMC infectious diseases. 2015 Dec;15(1):521. doi: 10.1186/s12879-015-1246-8

7. Jeena P, Thompson E, Nchabeleng M, Sturm A. Emergence of multi-drug-resistant *Acinetobacter anitratus* species in neonatal and paediatric intensive care units in a developing country: concern about antimicrobial policies. Annals of tropical paediatrics. 2001 Sep 1;21(3):245–51. https://doi.org/10.1080/02724930120077835

8. Gulen TA, Guner R, Celikbilek N, Keske S, Tasyaran M. Clinical importance and cost of bacteremia caused by nosocomial multi drug resistant *Acinetobacter baumannii*. International Journal of Infectious Diseases. 2015 Sep 1;38:32–5. doi: 10.1016/j.ijid.2015.06.014

9. Rumbo C, Gato E, López M, de Alegría CR, Fernández-Cuenca F, Martínez-Martínez L, Vila J, Pachón J, Cisneros JM, Rodríguez-Baño J, Pascual A. Contribution of efflux pumps, porins, and β-lactamases to multidrug resistance in clinical isolates of *Acinetobacter baumannii*. Antimicrobial agents and chemotherapy. 2013 Nov 1;57(11):5247–57. doi: 10.1128/AAC.00730-13

10. Manchanda V, Sanchaita S, Singh N. Multi-drug Resistant Acinetobacter. Journal of Global Infectious Diseases. 2010;2(3):291–304.

11. Heritier C, Poirel L, Nordmann P. Cephalosporinase over-expression resulting from insertion of ISAba1 in *Acinetobacter baumannii*. Clinical microbiology and infection. 2006 Feb 1;12(2):123–30. doi: 10.1111/j.1469-0691.2005.01320.x

12. Turton JF, Woodford N, Glover J, Yarde S, Kaufmann ME, Pitt TL. Identification of *Acinetobacter baumannii* by detection of the blaOXA-51-like carbapenemase gene intrinsic to this species. Journal of clinical microbiology. 2006 Aug 1;44(8):2974–6. doi: 10.1128/JCM.01021-06

13. Poirel L, Naas T, Nordmann P. Diversity, epidemiology, and genetics of class D β-lactamases. Antimicrobial agents and chemotherapy. 2010 Jan 1;54(1):24–38. doi: 10.1128/AAC.01512-08

14. Olive DM, Bean P. Principles and applications of methods for DNA-based typing of microbial organisms. Journal of clinical microbiology. 1999 Jun 1;37(6):1661–9.

15. Santimaleeworagun W, Thathong A, Samret W, Preechachuawong P, Sae-lim W, Jitwasinkul T. Identification and characterization of carbapenemase genes in clinical isolates of carbapenem-resistant *Acinetobacter baumannii* from a general hospital in thailand. Southeast Asian Journal of Tropical Medicine and Public Health. 2014 Jul 1;45(4):874.

16. Hornsey M, Loman N, Wareham DW, Ellington MJ, Pallen MJ, Turton JF, Underwood A, Gaulton T, Thomas CP, Doumith M, Livermore DM. Whole-genome comparison of two *Acinetobacter baumannii* isolates from a single patient, where resistance developed during tigecycline therapy. Journal of antimicrobial chemotherapy. 2011 May 12;66(7):1499–503. doi: 10.1093/jac/dkr168

17. Lowe M, Ehlers MM, Ismail F, Peirano G, Becker PJ, Pitout JD, Kock MM. *Acinetobacter baumannii*: epidemiological and beta-lactamase data from two tertiary academic hospitals in Tshwane, South Africa. Frontiers in Microbiology. 2018;9:1280. doi: 10.3389/fmicb.2018.01280

18. Anderson SE, Sherman EX, Weiss DS, Rather PN. Aminoglycoside Heteroresistance in *Acinetobacter baumannii* AB5075. mSphere. 2018 Aug 29;3(4):e00271–18. doi: 10.1128/mSphere.00271-18

19. Liakopoulos A, Miriagou V, Katsifas EA, Karagouni AD, Daikos GL, Tzouvelekis LS, Petinaki E. Identification of OXA-23-producing *Acinetobacter baumannii* in Greece, 2010 to 2011. Eurosurveillance. 2012 Mar 15;17(11):20117. https://doi.org/10.2807/ese.17.11.20117-en

20. Corrêa LL, Botelho LA, Barbosa LC, Mattos CS, Carballido JM, Castro CL, Mondino PJ, Paula GR, Mondino SS, Mendonça-Souza CR. Detection of blaOXA-23 in *Acinetobacter spp*. isolated from patients of a university hospital. Brazilian Journal of Infectious Diseases. 2012 Dec;16(6):521–6. doi: 10.1016/j.bjid.2012.10.003

21. Evans BA, Amyes SG. OXA β-lactamases. Clinical microbiology reviews. 2014 Apr 1;27(2):241–63. doi: 10.1128/CMR.00117-13

22. Perez F, Hujer AM, Hujer KM, Decker BK, Rather PN, Bonomo RA. Global challenge of multidrug-resistant Acinetobacter baumannii. Antimicrobial agents and chemotherapy. 2007 Oct 1;51(10):3471–84. doi: 10.1128/AAC.01464-06

23. Zarrilli R, Pournaras S, Giannouli M, Tsakris A. Global evolution of multi-drug-resistant *Acinetobacter baumannii* clonal lineages. Int J Antimicrob Agents. 2013 Jan 1;41(1):11–9. doi: 10.1016/j.ijantimicag.2012.09.008

24. Benmahmod AB, Said HS, Ibrahim RH. Prevalence and Mechanisms of Carbapenem Resistance Among *Acinetobacter baumannii* Clinical Isolates in Egypt. Microb Drug Resist. 2019 May 1;25(4):480–8. doi: 10.1089/mdr.2018.0141

25. Fournier PE, Richet H. The epidemiology and control of *Acinetobacter baumannii* in health care facilities. Clin Infect Dis. 2006 Mar 1;42(5):692–9. doi: 10.1086/500202

26. Cisneros JM, Rodríguez-Baño J. Nosocomial bacteremia due to *Acinetobacter baumannii*: epidemiology, clinical features and treatment. Clin Microbiol Infect. 2002 Nov 1;8(11):687–93. https://doi.org/10.1046/j.1469-0691.2002.00487.x

27. Chen TL, Wu RC, Shaio MF, Fung CP, Cho WL. Acquisition of a plasmid-borne blaOXA-58 gene with an upstream IS1008 insertion conferring a high level of carbapenem resistance to *Acinetobacter baumannii*. Antimicrobial agents and chemotherapy. 2008 Jul 1;52(7):2573–80. doi: 10.1128/AAC.00393-08

28. Qi C, Malczynski M, Parker M, Scheetz MH. Characterization of genetic diversity of carbapenem-resistant *Acinetobacter baumannii* clinical strains collected from 2004 to 2007. Journal of clinical microbiology. 2008 Mar 1;46(3):1106–9. doi: 10.1128/JCM.01877-07

29. Lowman W, Sriruttan C, Nana T, Bosman N, Duse A, Venturas J, Clay C, Coetzee J. NDM-1 has arrived: first report of a carbapenem resistance mechanism in South Africa. SAMJ: South African Medical Journal. 2011 Dec;101(12):873–5.

30. Chibabhai V, Perovic O. Epidemiology of carbapenem resistant Enterobacteriaceae at Charlotte Maxeke Johannesburg Academic Hospital. International Journal of Infectious Diseases. 2014 Apr 1;21:410. doi: https://doi.org/10.1016/j.ijid.2014.03.1265

31. Osei Sekyere J. Current state of resistance to antibiotics of last-resort in South Africa: a review from a public health perspective. Frontiers in public health. 2016 Sep 30;4:209. doi:10.3389/fpubh.2016.00209

32. Perovic O, Britz E, Chetty V, Singh-Moodley A. Molecular detection of carbapenemase-producing genes in referral Enterobacteriaceae in South Africa: A short report. South African Medical Journal. 2016;106(10):975–7. doi: 10.7196/SAMJ.2016.v106i10.11300.

33. Weber DJ, Rutala WA, Miller MB, Huslage K, Sickbert-Bennett E. Role of hospital surfaces in the transmission of emerging health care-associated pathogens: norovirus, Clostridium difficile, and Acinetobacter species. Am J Infect Control. 2010 Jun 1;38(5 Suppl 1):S25–33. doi: 10.1016/j.ajic.2010.04.196

34. Nordmann P, Poirel L. *Acinetobacter baumannii*: basic and emerging mechanisms of resistance. Eur Infect Dis. 2008;2:94–97.

35. Coyne S, Courvalin P, Périchon B. Efflux-mediated antibiotic resistance in *Acinetobacter spp.* Antimicrob Agents Chemother. 2011 Mar 1;55(3):947–53. doi: 10.1128/AAC.01388-10

36. Marchand I, Damier-Piolle L, Courvalin P, Lambert T. Expression of the RND-type efflux pump AdeABC in *Acinetobacter baumannii* is regulated by the AdeRS two-component system. Antimicrob Agents Chemother. 2004 Sep 1;48(9):3298–304. doi: 10.1128/AAC.48.9.3298-3304

37. Lari AR, Ardebili A, Hashemi A. AdeR-AdeS mutations & overexpression of the AdeABC efflux system in ciprofloxacin-resistant *Acinetobacter baumannii* clinical isolates. The Indian journal of medical research. 2018 Apr 1;147(4):413. doi: 10.4103/ijmr.IJMR_644_16

38. Higgins PG, Janssen K, Fresen MM, Wisplinghoff H, Seifert H. Molecular epidemiology of *Acinetobacter baumannii* bloodstream isolates obtained in the United States from 1995 to 2004 using rep-PCR and multilocus sequence typing. J Clin Microbiol. 2012 Nov 1;50(11):3493–500. doi: 10.1128/JCM.01759-12

39. Bratu S, Landman D, Martin DA, Georgescu C, Quale J. Correlation of antimicrobial resistance with beta-lactamases, the OmpA-like porin, and efflux pumps in clinical isolates of *Acinetobacter baumannii* endemic to New York City. Antimicrob Agents Chemother. 2008 Sep 1;52(9):2999–3005. doi: 10.1128/AAC.01684-07

40. Antunes L, Visca P, Towner KJ. *Acinetobacter baumannii*: evolution of a global pathogen. Pathogens and disease. 2014 Aug 1;71(3):292–301. doi: 10.1111/2049-632X.12125

41. Lean SS, Suhaili Z, Ismail S, Rahman NI, Othman N, Abdullah FH, Jusoh Z, Yeo CC, Thong KL. Prevalence and genetic characterization of carbapenem-and polymyxin-resistant *Acinetobacter baumannii* isolated from a tertiary hospital in Terengganu, Malaysia. ISRN microbiology. 2014 Mar 19;2014. doi: 10.1155/2014/953417

42. Jasovský D, Littmann J, Zorzet A, Cars O. Antimicrobial resistance—a threat to the world’s sustainable development. Upsala journal of medical sciences. 2016 Jul 2;121(3):159–64. doi: 10.1080/03009734.2016.1195900

43. Sherertz RJ, Sullivan ML. An outbreak of infections with *Acinetobacter calcoaceticus* in burn patients: contamination of patients’ mattresses. Journal of Infectious Diseases. 1985 Feb 1;151(2):252–8. doi: 10.1093/infdis/151.2.252

44. Choi WS, Kim SH, Jeon EG, Son MH, Yoon YK, Kim JY, Kim MJ, Sohn JW, Kim MJ, Park DW. Nosocomial outbreak of carbapenem-resistant *Acinetobacter baumannii* in intensive care units and successful outbreak control program. Journal of Korean medical science. 2010 Jul 1;25(7):999–1004. doi: 10.3346/jkms.2010.25.7.999

45. Voets GM, Fluit AC, Scharringa J, Stuart JC, Leverstein-van Hall MA. A set of multiplex PCRs for genotypic detection of extended-spectrum β-lactamases, carbapenemases, plasmid-mediated AmpC β-lactamases and OXA β-lactamases. International journal of antimicrobial agents. 2011 Apr 1;37(4):356–9. doi:10.1016/j.ijantimicag.2011.01.005

46. Dallenne C, Da Costa A, Decré D, Favier C, Arlet G. Development of a set of multiplex PCR assays for the detection of genes encoding important β-lactamases in Enterobacteriaceae. Journal of Antimicrobial Chemotherapy. 2010 Jan 12;65(3):490–5. doi: 10.1093/jac/dkp498

47. Beheshti M, Talebi M, Ardebili A, Bahador A, Lari AR. Detection of AdeABC efflux pump genes in tetracycline-resistant Acinetobacter baumannii isolates from burn and ventilator-associated pneumonia patients. Journal of pharmacy & bioallied sciences. 2014 Oct;6(4):229. doi: 10.4103/0975-7406.142949

